# Beyond the CTCT: Remote Structural Changes After VIM-MRgFUS in Essential Tremor

**DOI:** 10.1101/2024.10.08.617157

**Authors:** Jonas Krauß, Neeraj Upadhyay, Veronika Purrer, Valeri Borger, Marcel Daamen, Angelika Maurer, Carsten Schmeel, Alexander Radbruch, Ullrich Wüllner, Henning Boecker

## Abstract

**Introduction:** Essential tremor (ET) is a progressive disorder characterized by altered network connectivity between the cerebellum, thalamus, and cortical regions. Magnetic Resonance-guided Focused Ultrasound (MRgFUS) of the ventral intermediate nucleus (VIM) is an effective, minimally invasive treatment for ET. Increasing clinical use in ET facilitates research on structural changes after thalamic lesions.

**Methods:** Forty-six patients with medication-resistant ET underwent unilateral VIM-MRgFUS. Voxel-based morphometry was applied to investigate Gray Matter Volume (GMV) changes over a time span of 6 months in the whole brain and the thalamus in particular to investigate local and distant effects.

**Results:** Clinically, contralateral tremor significantly decreased by 68 % at 6 months following MRgFUS. In addition to local changes in thalamic nuclei (VIM, ventral lateral posterior, centromedian thalamus and pulvinar), VBM revealed remote GMV decreases in the ipsilesional insula and the anterior cingulate cortex as well as the contralesional middle occipital gyrus. Increased GMV was found in both temporal gyri. There was no significant correlation between regional GMV declines and tremor improvement. However, temporal volume increases were associated with improved motor-related functional abilities and quality of life outcomes.

**Conclusion:** Our findings implicate distributed structural changes following unilateral VIM-MRgFUS. Structural losses could reflect Wallerian degeneration of VIM output neurons or plasticity due to decreased sensory input following tremor improvement.

## Introduction

Essential tremor (ET) is a slowly progressing neurological disorder widely recognized as a network pathology. Despite extensive research, its pathophysiology remains incompletely understood^1^. Neuroimaging studies using functional magnetic resonance imaging (fMRI) and positron emission tomography (PET) have highlighted alterations in functional connectivity within and between the cerebellum, thalamus, and various cortical motor regions as key contributors to ET pathogenesis^2,3^. Dysfunctions in aberrant cerebellar-thalamo-cortical loops are believed to play a pivotal role in the generation and propagation of tremor oscillations. Notably, the cerebellar dentate nucleus and the primary motor cortex (M1) are connected via the thalamic ventral intermediate nucleus (VIM) through the cerebello-thalamo-cortical tract (CTCT) which is implicated in ET pathophysiology^4^.

Pharmacotherapy using β-blockers or primidone is the first-line therapy, but effectively controls symptoms in only approximately 50% of patients and may lead to intolerable side effects for many patients^5,6^. As an alternative, stereotactic treatments have become well-established in recent years. Deep brain stimulation (DBS) is the most common surgical intervention, involving unilateral or bilateral implantation of electrodes in key areas of the pathological network (for ET, typically: VIM) to modulate aberrant neural activity via electrical currents. In addition to DBS, Magnetic Resonance-guided Focused Ultrasound (MRgFUS) offers an incisionless procedure that may surpass other thalamotomy procedures such as radio-surgical gamma knife^6^. Long-term treatment success has been demonstrated^7^ but the adaptive processes associated with MRgFUS lesioning, need further investigation.

Using probabilistic tractography, we previously^8^ reported significant reductions in streamline counts within the CTCT and a decrease in excitatory input from the VIM to the ipsilateral M1 following VIM MRgFUS treatment in ET patients. These findings align with a recent study^9^, which described functional reorganization within the CTCT network encompassing M1, but also extending beyond, to dorsal attention (anterior cingulate cortex, superior temporal gyrus), and visual areas.

While numerous studies have documented functional changes following VIM-MRgFUS^8,10,11^, studies on subsequent gray matter volume (GMV) change remain sparse, particularly at the whole-brain level. Observational patient studies have linked GMV changes to motor and non-motor symptoms in both PD^12^ and ET^13^, while treatment studies in ET focusing on DBS have investigated morphological changes primarily in brain areas connected via the CTCT^14^. In addition, GMV changes associated with tremor reduction have been described in PD patients following VIM-MRgFUS treatment^15^. The structural reorganization upon MRgFUS in ET is somewhat unexplored. Remote network changes might explain further facets of clinical improvement and potential side effects of VIM lesioning using MRgFUS.

In this study we analyzed GMV changes at the whole-brain level using voxel-based morphometry (VBM), along with region-of-interest (ROI) analysis of thalamic subnuclei to further elucidate MRgFUS-related local structural adaptation processes. In particular, we expected to find neuroplastic changes in interconnected areas and thus tried to establish relationships between the exact location and size of the lesion and the corresponding morphologic changes, as well as their impact on clinical improvement. We hypothesized GMV decreases in brain structures linked directly via the CTCT, but also explored remote areas anatomically connected to the lesion center.

## Methods

### Patients

Fifty-two patients with diagnosis of definitive drug-refractory ET were recruited from the University Hospital Bonn, Germany. All patients underwent unilateral VIM-MRgFUS (ExAblate Neuro 4000, Insightec) between 2018 and 2023. Tremor presented bilaterally in every patient. Treatment targeted either the more affected hand or the dominant hand. Inclusion criteria are listed in the German Clinical Trials Registry (DRKS00016695). Apart from MRI-specific factors (e.g., claustrophobia, implanted electric devices, etc.), further exclusions criteria were brain scan irregularities, and comorbid psychiatric or neurological conditions. After excluding six patients due to poor image quality, the final sample included 46 patients (12 female, age: 73.41 ± 7.42 (mean ± standard deviation)). Seven patients were treated on the right VIM, and 39 on the left.

The study was approved by the local ethics board of the University Hospital Bonn (No. 314/18), with informed consent obtained in accordance with the Declaration of Helsinki

### Study design

In this pre-post study design, MRI examinations were performed two days before the MRgFUS procedure, 3 days post, and 6 months post procedure. Additionally, clinical scales were administered to monitor patient-related outcomes of the MRgFUS procedure.

### Clinical assessment

To measure long-term clinical outcome, tremor assessments using the Fahn-Tolosa-Marin Clinical Rating Scale for Tremor (FTM)^16^ before and 6 months after MRgFUS were compared, for the treated and untreated side separately. For this analysis, only upper extremity scores were included. Higher FTM scores indicate a more severe tremor presentation. The FTM comprises three parts: Part A + B which assess tremor severity and was summed up as FTM-A/B for the analysis and Part C which examines the impairments in activities of daily living (ADL).

Additionally, subjects completed the 36-Item Short Form Survey Instrument^17^ (SF-36), a questionnaire measuring eight dimensions of health-related quality of life (QoL). Higher scores indicate better QoL per domain. Both tests are further described in Supplementary Table 1.

### MRgFUS procedure

Details of the MRgFUS procedure were described previously^18^. Briefly, before MRgFUS, a clinical CT was performed to assess contraindications and confirm a skull density ratio of ≥0.3. For all but the first three patients, lesion site planning utilized preoperative probabilistic tractography from diffusion tensor imaging to map sensorimotor pathways (e.g., corticospinal tract, DRT, medial lemniscus). During the procedure, subthreshold sonications (<50 °C) were applied to assess tremor reduction and monitor side effects. Final treatment sonications (>55 °C) were delivered to the site showing optimal tremor reduction without adverse effects.

### MRI data acquisition

All study participants underwent brain imaging with a 3T MRI scanner (Achieva TX, Philips Healthcare, Best, The Netherlands) equipped with a 8-channel head coil 2 days before, 3 days after, and 6-months after MRgFUS. A T1-weighted three-dimensional magnetization-prepared rapid gradient-echo (MPRAGE) sequence was acquired to obtain high-quality isotropic anatomical images. The following scanning parameters were used: 3D-FFE sequence; matrix size: 256 × 256 mm; TE/TR: 3.9/7.3 ms; flip angle: 15°; spatial resolution (acquired voxel size): 1 × 1 × 1 mm^3^ with 0 mm interslice gap over 180 slices; scan duration: 4:40 min.

### Manual Lesion segmentation

Lesion segmentation was in accordance with previously reported procedures^19^. In native space, the lesion was segmented using FSLeyes (Version 0.34.2). The maps were then nonlinearly normalized to the 1.5 mm voxel-size MNI template (International Consortium for Brain Mapping 2009b nonlinear, asymmetric; Montreal Neurological Institute, Montreal, Canada) using Advanced Normalization Tools (ANTs, Philadelphia, PA, Version 2.3.1). Normalized lesion maps from both time points were summed up to identify areas with highest lesion/edema overlap among patients.

### Image preprocessing and Voxel-Based Morphometry (VBM)

The VBM analysis only compared images at baseline and 6 months following MRgFUS. After converting the DICOM image series to Nifti format, they were visually checked for artifacts and the origin manually set to the anterior commissure. Images of patients operated on the right VIM were flipped using the swap dimension command in FSL (FSL6.0.4, http://www.fmrib.ox.ac.uk/fsl). VBM was conducted using the longitudinal pipeline of the Computational Anatomy Toolbox 12 (CAT12.8.2, University Hospital Jena, Jena, Germany)^20^ for Statistical Parametric Mapping 12 (SPM12, Wellcome Department of Cognitive Neurology, London, UK). SPM12 and CAT12 were running in MATLAB R2022a (The MathWorks Inc., Natick, MA, USA). Preprocessing involved segmenting the images into gray matter (GM), white matter, and cerebrospinal fluid probability maps. Tissue segmentations were registered to the MNI152NLin2009cAsym template using Diffeomorphic Anatomical Registration Through Exponentiated Lie algebra (DARTEL), with voxel size set to 1.5mm. GMV segmentations underwent visual inspection, followed by quality control using CAT12s built-in function, with scans exceeding a z-score of 1.0 excluded. Finally, GMV maps were smoothed using a 6mm full width, half maximum isotropic Gaussian kernel.

### Statistical method

Assumption testing and inferential statistical analysis of the demographic and clinical data was conducted with SPSS (Version 27, IBM Corp., 2020). Outliers were checked, and normality was confirmed by visually inspecting QQ-plots. Pairwise t-tests were used to assess differences between the two time points.

For statistical voxel-wise comparison of the MRIs at whole-brain level, a pairwise t-test (Baseline (BL) vs 6-month post (6m)) was conducted using SPM12. The clusters were obtained at voxel-wise *p*<0.001 and further corrected for multiple comparisons using cluster-level family-wise-error correction (p<0.05 FWEc) according to Gaussian Random Field theory. Clusters were labeled according to the Automated Anatomical Labeling atlas version 3 (AAL3)^21^.

Based on the lesion segmentation results, we further characterized the local thalamic effects of MRgFUS by a ROI-analysis, using the CAT12 preinstalled thalamic nuclei atlas by Saranthan et al. (2021)^22^ that parcellates the thalamus in 12 nuclei per hemisphere. To adequately correct for multiple comparisons for this smaller volume we applied a Bonferroni-Holm correction using *p* = 0.05.

To explore associations between VBM changes and clinical outcomes, mean beta values from significant clusters were correlated with tremor scores (FTM-A/B) and quality of life (QoL) changes (FTM-C, SF-36 subscores) using Pearson correlation coefficients. Change scores were calculated as the difference between 6 months and baseline, with these analyses considered exploratory and not corrected for multiple comparisons.

## Results

### Clinical outcomes

The FTM tremor scores decreased significantly (t(45) = 15.71, *p* < .001) from BL (M = 19.51, SD = 5.20) to 6m post-MRgFUS (M = 6.28, SD = 4.86) contralateral to the treated ViM, which reflects a mean improvement of ∼70 %. The non-treated side did not show a significant difference (t(45) = 0.63, *p* = .531) from BL (M = 17.47, SD = 6.10)) to 6m (M = 17.14, SD = 6.38) (see Figure 1).

**Figure 1.**
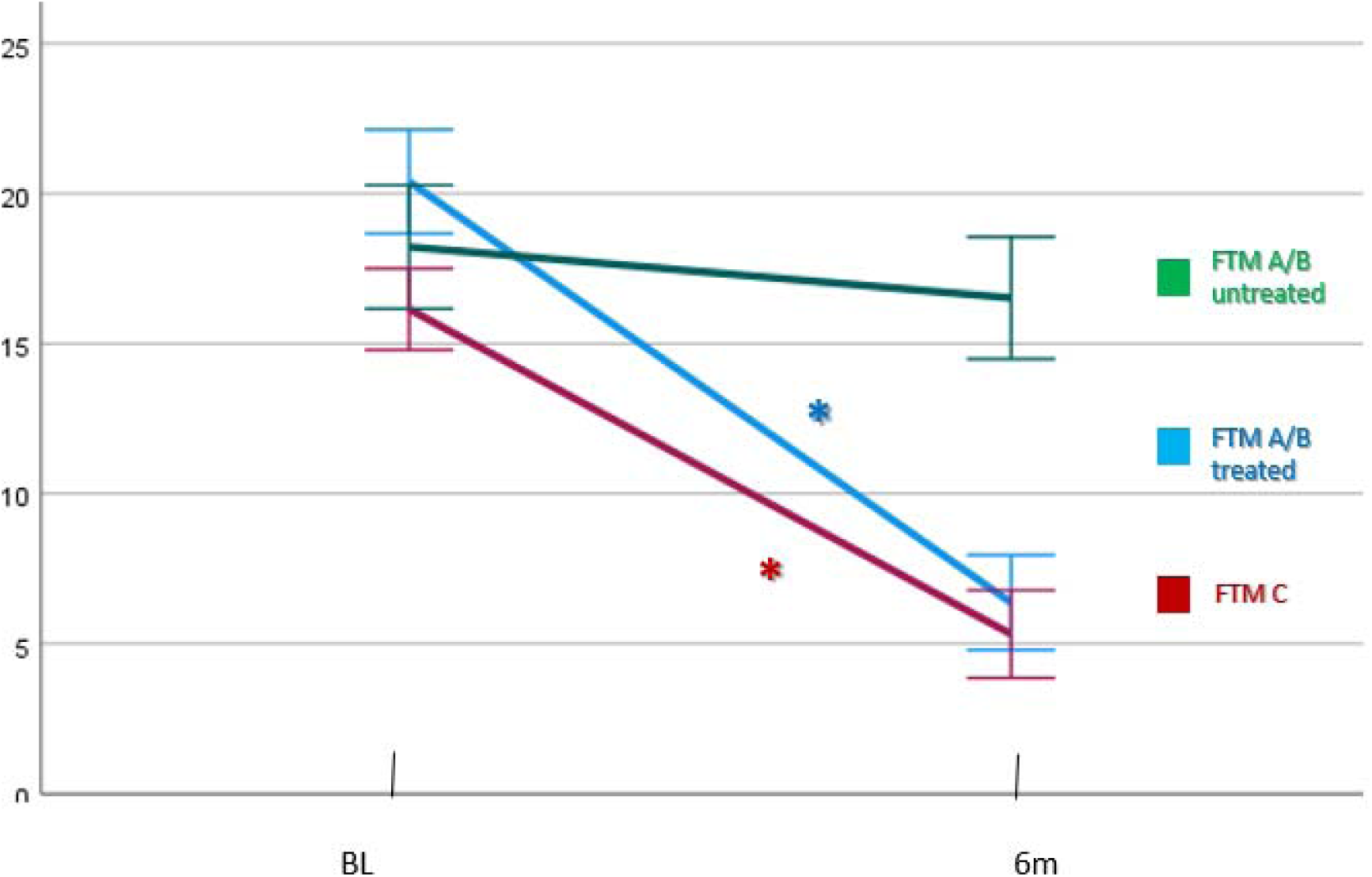
FTM changes T0 versus 6 month post MRgFUS. FTM scores from T0 compared to 6 months following VIM-MRgFUS. Significant changes are visualized with an asterisk (*). The error bars represent standard deviations. Subscore of FTM-A and FTM-B were summed (FTM A/B) to indicate general tremor severity.

ADL parameter (FTM part C) significantly improved by 66 % (t(45) = 14.52, *p* < .001) from BL (M = 15.76, SD = 4.11) to 6m (M = 5.38, SD = 4.66).

Comparing the scores at BL to 6m of the SF-36, the following QoL domains showed significant improvement: Physical and Emotional Role Functioning, Emotional Wellbeing, Social Functioning and General Health (Table 2).

### Lesion segmentation

Lesion and edema regions were segmented for all included patients at 3 days post-procedure (shown in red/yellow in Figure 2f) as well as in 34 patients 6 months-post procedure (shown in blue in Figure 2). Edema is known to increase in size shortly after MRgFUS^18^, gradually shrinking as recovery progresses. This suggests that the actual lesion is concentrated around the center of mass seen in the 3-day post-procedure heatmap (yellow, Figure 2f), which primarily aligns with the VIM area. The overlap of this lesion and edema region extends into adjacent areas, including the pulvinar (PV), centromedian (CM), and posterior ventrolateral thalamus (VLp).

**Figure 2.**
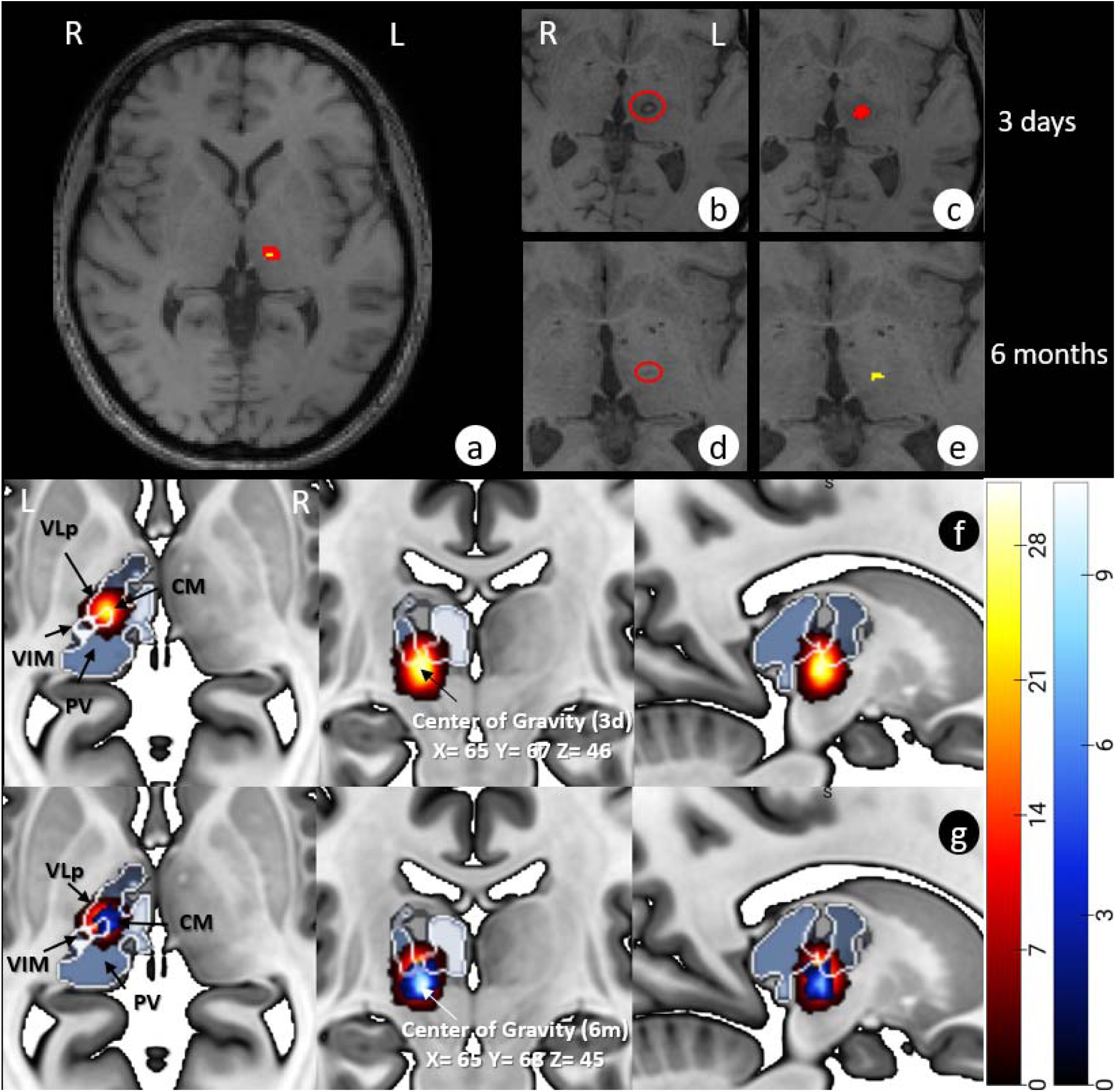
Lesion segmentation workflow and overlap of 3 days and 6 months following MRgFUS. (a) Lesion overlap in MNI-space showing volume reduction from 3 days (3d) to 6 months (6m). (b-c) Lesion and edema 3 days post-treatment. (d-e) Same at 6m, showing edema recession. (f-g) Combined lesion masks 3d (red/yellow) and 6m (blue/white).

Six months after procedure, the size of the hypointensities visible in the MPRAGE substantially decreased, leaving only a smaller lesion area localized to the VIM area, but still reaching the VLp and CM (Figure 2 d, e, g). Figure 2 provides a detailed illustration of this process and its outcomes.

### Thalamus ROI-based VBM analysis

To investigate regional changes in the thalamic nuclei in consideration of the lesion segmentation, we conducted an atlas-based ROI-analysis. Consistent with findings from the segmentation, the GMV decline extended beyond the target location (VIM) and encompassed reductions in the VLp, CM, and PV. Refer to Figure 2 for visual representation of these results.

### Whole-brain VBM analysis

Whole-brain structural changes from BL to 6m following VIM-MRgFUS revealed both local and remote alterations in GMV (Figure 4). As expected, the highest beta values were observed in the thalamus, particularly in the target lesion center (VIM) and surrounding thalamic nuclei. Additionally, GMV decline was detected in left insula, Rolandic operculum, left anterior cingulate cortex (ACC), and right middle occipital gyrus. Conversely, increases in GMV were observed in the right superior and middle temporal gyrus, as well as in the left inferior temporal gyrus.

With a more lenient threshold (p = 0.001, uncorrected), additional trends to decreases at peak-level were found in the left middle occipital gyrus, the right temporal gyrus as well as the right cerebellar lobule VIII. See Supplementary Table 2 and 3 for the statistics of each GMV decrease and increase at cluster-level.

### Correlation with clinical data

To assess associations between structural changes and clinical scores, we correlated the mean beta values from each significant cluster of the voxel-wise analysis with FTM-A/B (tremor severity) as well as FTM-C (ADL), and the SF-36 subscales (indicating changes in QoL domains).

The reductions of the FTM-C showed a significant inverse correlation with GMV increase in the left inferior temporal gyrus (r = -0.434, p = 0.007). In addition, improvement in the SF-36 domain Physical Role Limitation (r = 0.447, p = 0.015) and Physical Functioning (r = 0.368, p = 0.050) showed correlations with GMV increase in right middle temporal gyrus.

Emotional Wellbeing was associated with GMV increase in the right middle temporal lobe (r = 0.506, p = 0.005). Moreover, there was a positive correlation between Social Functioning (r = 0.380, p = 0.042) and left middle temporal lobe increase as well as between improvement in General Health and left middle temporal lobe increase (r = 0.537, p = 0.003).

## Discussion

Our study revealed significant GMV changes following MRgFUS VIM-lesioning which induced substantial contralateral tremor reduction. We observed GMV decreases in the lesioned VIM but also in the PV, the VLp and the CM. Additionally, remote GMV decreases were noted in the ipsilateral insula, the ipsilateral ACC, and the contralesional middle occipital gyrus (as well as at trend levels in the ipsilesional middle occipital gyrus). On the other hand, we found increases in GMV in bilateral temporal gyri. Findings suggest complex neuroanatomical changes post-treatment, reflecting both direct effects of ablation and broader network-level adaptations. While tremor improvement did not correlate with thalamic GMV decrease, improvements of functional disabilities, emotional well-being, social functioning and general health were significantly correlated with GMV increases in bilateral temporal areas.

At three days post-MRgFUS, MR images showed signal alterations extending beyond the ablation target, reflecting both the lesion and surrounding edema (Figure 2a-b). Group-level density maps of these alterations (Figure 2f) revealed the involvement of the VIM and surrounding nuclei (VLp, CM, PV). At six months, focal signal alterations were visible in about 75% of patients (Figure 2g), with segmentation overlap still extending beyond the VIM to include the VLp and CM, indicating interindividual variability in tissue damage. VBM analysis confirmed GMV reductions in the VIM, VLp, and CM, while also detecting PV changes. These findings highlight that MRgFUS primarily affects the VIM but also impacts adjacent thalamic nuclei. The lack of visible lesions in the PV at six months (Figure 2g) alongside significant VBM changes (Figure 3) suggests protracted structural changes, possibly triggered by initial edema and presumably also inflammatory responses.

**Figure 3.**
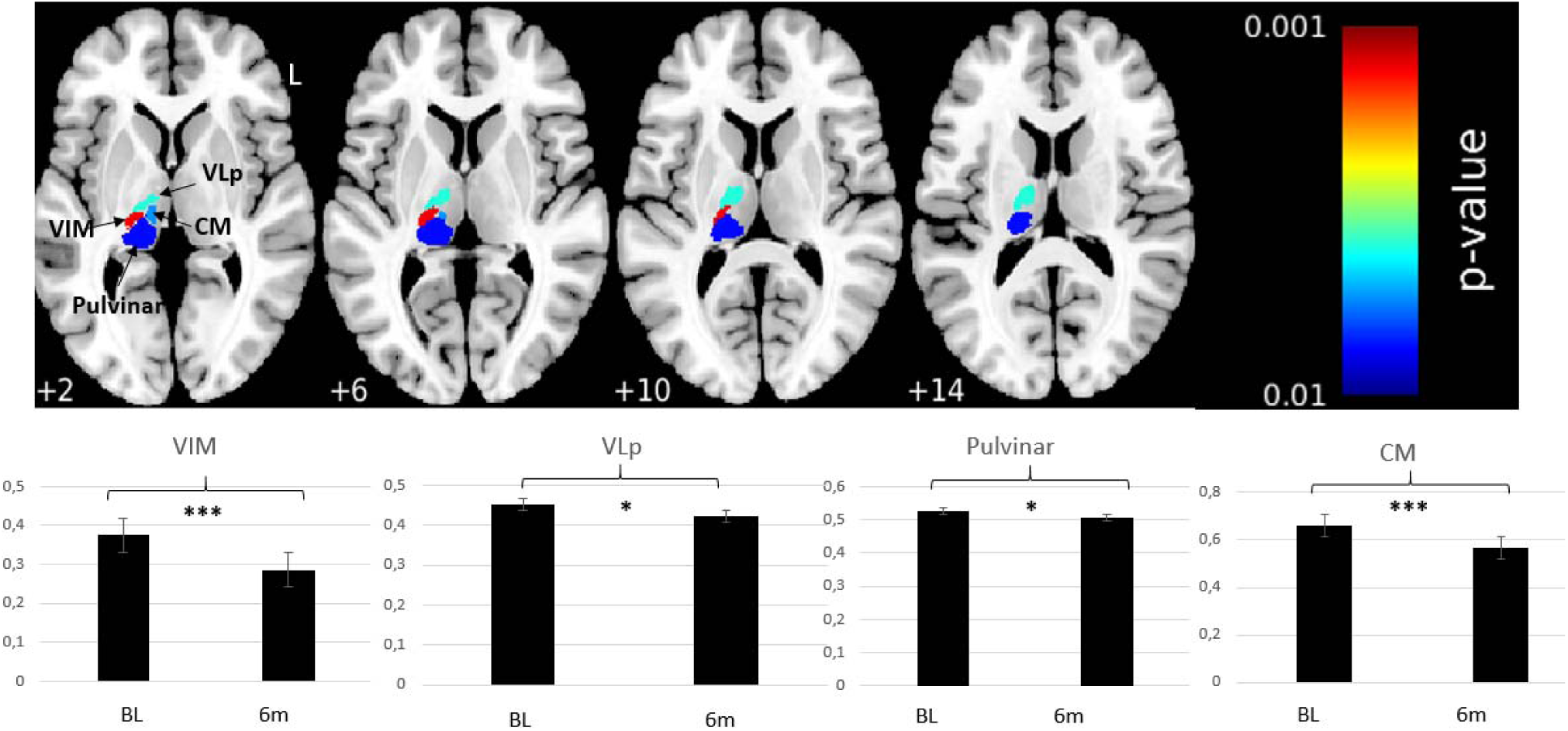
ROI-based statistical analysis of thalamic subregions: 6 months post MRgFUS versus baseline. Significant GMV reductions at 6m in VIM, CM, VLp, and PV using atlas^22^. Multiple corrections with Bonferroni-Holm (FWE, p = .05). Bar charts show GMV at baseline and 6m with standard error (*p = .05, ***p < .001).

**Figure 4.**
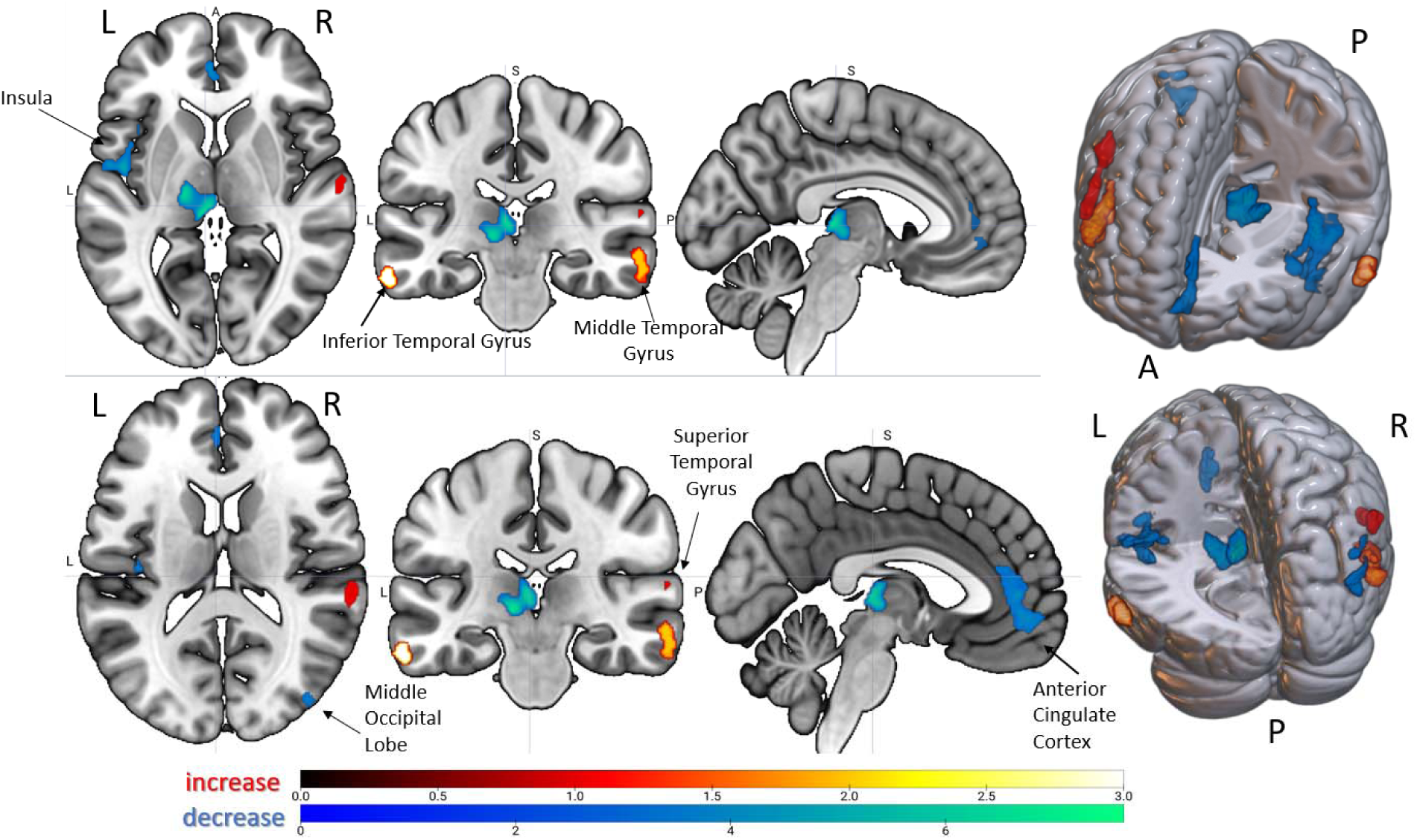
Whole-brain statistical analysis of gray matter changes: 6 months post MRgFUS versus baseline. Significant GMV changes in left thalamus, insula/Rolandic operculum, ACC, right middle occipital gyrus (decreases), and temporal gyri (increases). Multiple comparison correction method: FWE cluster-level correction (Decrease: k=239, Increase: k=290). Color bars show t-values: green (GMV increase), red/yellow (GMV decrease).

**Table 1.** Clinical score changes following VIM-MRgFUS.

| Clinical test/domain | Mean ± SD<br>BL | Mean ± SD<br>6m | t-value | p |
| --- | --- | --- | --- | --- |
| **FTM** |  |  |  |  |
| A&B (treated side) | 19.51 (±5.20) | 6.28 (±4.86) | 15.71 | <.001* |
| A&B (untreated side) | 17.47 (±6.10) | 17.14 (±6.34) | 0.63 | .531 |
| C | 15.76 (±4.11) | 5.38 (±4.66) | 14.52 | <.001* |
| **SF-36** |  |  |  |  |
| Physical Functioning | 71.03 (±28.79) | 75.86 (±25.95) | -0.93 | .361 |
| Role Limitation (Physical) | 48.27 (±41.69) | 65.52 (±40.87) | -2.62 | .014* |
| Role Limitation (Emotional) | 64.37 (±46.23) | 85.06 (±32.84) | -2.31 | .029* |
| Vitality | 50.00 (±25.46) | 55.35 (±22.83) | -1.07 | .295 |
| Emotional Wellbeing | 56.28 (±25.21) | 73.38 (±20.21) | -3.76 | .001* |
| Social Functioning | 67.50 (±36.15) | 82.93 (±24.12) | -2.92 | .007* |
| Bodily Pain | 71.37 (±31.05) | 74.66 (±28.47) | -0.59 | .559 |
| General Health | 44.59 (±20.20) | 58.24 (±25.26) | -2.99 | .006* |

*Note.* Results of pairwise t-tests comparing BL to 6m. Significant p-values are marked with an asterisk (\*). FTM-A/B is the sum for FTM-A and FTM-B of the upper extremities used to indicate tremor severity. BL=Baseline, 6m=6-months follow-up, SD=Standard deviation

Whole-brain analysis at six months also indicates remote structural alterations (GMV decreases and increases) after the MRgFUS intervention, suggesting that lesion-induced changes are not confined to the CTCT but extend to distant yet connected cortical regions. As commonly seen in stroke patients, GMV changes can be caused by deafferentation of remote brain structures connected with a lesion site. This process, known as Wallerian degeneration, has also been proposed as an underlying mechanism in the MRgFUS literature^19,23^. We observed a GMV decrease in the ipsilateral insula and ACC. While a precise map of thalamic subnuclei connections to the insula and ACC in humans has not yet been established, the thalamus, insula, and ACC are known to be structurally connected and form networks associated with a range of sensory and cognitive functions^24^.

Non-human primate studies using anterograde and retrograde tracing have highlighted connections of thalamic nuclei, located ventrally and within the intralaminar group (e.g., CM) and the pulvinar^25–27^. Correspondingly, DTI analyses in humans demonstrated that the intralaminar group and specifically the medially located nuclei (e.g., CM) are connected to various cortical structures including the insula, suggesting involvement in somatosensory, visual, olfactory, and gustatory processing^28^. Given that we observed significant GMV decline in ventral subnuclei near the lesion, as well as in the intralaminar group, the GMV decrease in the insula is likely mediated through Wallerian degeneration.

For the ACC, animal studies^29^ suggest strong structural connections from the thalamus, particularly from ventral and medial areas. While these connections might exist in humans too^30,31^, subnuclei data are inchoate and to date primarily show structural connections from the mediodorsal thalamus to the ACC^32^. Indeed, our data suggested some GMV decrease in the mediodorsal thalamus, but this change was not significant.

Functionally, the insula and ACC are integral components of the salience network, which integrates autonomic and behavioral responses to external stimuli influencing motor regulation, with the thalamus acting as a control hub^24,33^. Supporting this, Fang and colleagues^34^ reported increased functional activity in the salience network among ET patients. Additionally, ET patients often show GMV increases in the frontal lobe and left insula, indicating heightened functional load in these regions^35^. The findings of GMV declines in the left insula and ACC after treatment, might also be attributable to reduced functional demands following tremor improvement, leading to adaptive structural changes in these areas. Diminution of tremor likely reduces the sensory and attentional system requirements, potentially resulting in structural changes as these areas are no longer over-engaged. Taylor and colleagues^36^ proposed a resting-state connectivity network between the insula and the cingulate cortex that is involved in skeletomotor body orientation and monitoring. Considering these roles, activity in this network may be reduced due to the absence of tremor which would decrease the demand for motor control and possibly induce intermediate structural reorganization.

Also, the GMV decline in the contralesional middle occipital gyrus might be related to altered sensory input. ET patients showed GMV increases relative to healthy controls in visual areas, suggesting increased demands for visuospatial control in the presence of intention tremor^37^. Moreover, structural and functional changes have been observed frequently in visual areas following tremor reduction due to VIM lesioning or DBS^11,15,38^. These adaptations may be caused by normalization of the interconnectivity of visual areas due to tremor improvement^39^. A significant decrease was observed primarily contralateral to the lesion, but the anatomically connected ipsilateral primary visual cortex also showed a GMV decline at a more lenient threshold.

Our finding of increased GMV in bilateral temporal areas aligns with observations in untreated ET patients showing decreased GMV in comparison with healthy controls^40^. Previous neuroimaging and electrophysiological recording studies^41,42^ suggested that temporal areas are involved in control of goal-directed movement sequences and sensorimotor coordination. After diminution of tremor, the increase in motor activities involving fine hand movements (e.g., using cutlery, brushing teeth, etc.) may have influenced the demand for sensorimotor control and triggered GMV increases in temporal regions. Supporting this, a DBS connectomics study^43^ highlighted the functional relationship between temporal areas, along with visual and motor regions, and tremor improvement. Likewise, our study found a correlation between increased temporal GMV and improved functional abilities in social life and daily living, as measured by the FTM-C and the SF-36. It has to be acknowledged, however, that other studies report contrary results, namely increased GMV in the superior and middle temporal areas when comparing ET patients with healthy controls^37^.

While our findings show widespread cortical GMV changes predominantly ipsilateral to the MRgFUS lesion, it is surprising that no significant changes were observed in M1, the primary output region of the VIM. Previous DTI data from our group^8^ highlighted reductions in streamline counts within the CTCC, particularly in fibers connecting the VIM to ipsilateral M1. Complementing this, task-based fMRI revealed decreased activation in the ipsilateral M1^8^. However, the literature on structural changes remains inconclusive: Bolton et al.^44^ observed structural decreases in the precentral gyrus and insula after Gamma Knife VIM-ablation, while, conversely, Bruno and colleagues^23^ found volumetric declines in the putamen, pallidum, and cerebellar cortex following MRgFUS in a small sample of ET patients but did not detect changes in the motor cortex. The variability in findings may depend on study-specific factors, such as follow-up intervals, and further research is needed to clarify structural alterations in VIM-output regions. Similar to the aforementioned morphometric studies^23,44^, no association between tremor improvement (FTM-A/B) and GMV changes were found in our study. While this is somewhat unexpected for the thalamic lesion site, the lack of a direct association for regions not involved in tremor generation is more plausible.

Limitations are as follows: This study did not include healthy controls for comparison which could have clarified whether the structural alterations seen in ET patients reflect baseline abnormalities or shifts toward a healthier state. Longer follow-up intervals may reveal additional variability with regard to the stability of the tremor improvements^45,46^. Moreover, the inherent heterogeneity in both the clinical presentation of ET patients (e.g., the intensity and type of tremor) and the variability in VIM lesioning locations limits the generalizability of our findings, even with an adequate sample size. Other potential confounding variables like medication use, disease duration or side effects were not controlled for, which could influence the results.

## Conclusion

Our findings implicate local but also more distributed structural adaptations in areas beyond the CTCT network following unilateral VIM-MRgFUS. Structural losses could reflect Wallerian degeneration of VIM output neurons and adjacent thalamic subnuclei; or, alternatively, plasticity due to decreased sensory input following tremor improvement. While structural changes were not correlated with tremor decrease, consistent association of GMV increases in temporal areas with improved QoL were reported. In summary, these findings are significant for understanding neuroplasticity following tremor treatment in ET and motivate further research into biomarker of tremor improvement, side effects and shifts in functional demand.

## Relevant conflicts of interest/financial disclosures

The authors declare no conflicts of interest related to this study.

## Funding agency

The MRgFUS facility was funded by the German Research Foundation (INST 1172/64-1) and the Medical Faculty of Friedrich-Wilhelms-University Bonn.

## Acknowledgment

We are grateful to the patients for their participation in this study. Open Access funding enabled and organized by Project DEAL.

## Authors’ Roles

J.K. contributed to analysis, and interpretation; drafted the manuscript; and critically revised the manuscript.

N.U. contributed to analysis and interpretation, and critically revised the manuscript.

V.P. contributed to data acquisition and critically revised the manuscript

V.B. contributed to data acquisition and critically revised the manuscript

M.D. contributed to interpretation and critically revised the manuscript

A.M. critically revised the manuscript

A.R. critically revised the manuscript

U.W. contributed to data acquisition and critically revised the manuscript

H.B. contributed to analysis and interpretation, and critically revised the manuscript

All authors provided their final approval and are accountable for all aspects of the work in ensuring that questions related to the accuracy or integrity of any part of the work are appropriately investigated and resolved.

## Funding

The MRgFUS facility was funded by the German Research Foundation (INST 1172/64–1).

## Competing interests

U.W. has received consultancy fees from STADA Pharm, Idorsia and Zambon. He has received lecture fees from Abbvie, Bayer, Bial, Roche and Zambon. He has received study support from the DFG, the German Ministry of Research (BMBF), the International Parkinson

Fund and the German Parkinson Association (dPV), as well as from the German Center for Neurodegenerative Diseases (DZNE) and the University of Bonn.

The other authors declare that they have no known competing financial interests or personal relationships that could have appeared to influence the work reported in this paper.

## Supplementary material

**Supplementary Table 1.** Used clinical scales and descriptions.

Used clinical scales and descriptions
| **FTM** |  |
| --- | --- |
| **Domain** | **Description** |
| Part A | A quantitative visual rating of tremor severity in different body parts (e.g., face, upper extremities, lower extremities) with respect of tremor type (e.g. postural or intention tremor) |
| Part B | An evaluation of the patient's ability to perform tasks requiring steady line drawing and writing. |
| Part C | A subjective assessment of limitations in activities of daily living (ADL; e.g., eating, drinking, hygiene). |
| **SF-36** |  |
| **Domain** | **Description** |
| **Physical Functioning** | Limitations in physical activities due to health problems |
| **Social Functioning** | Limitations in social activities due to physical or emotional problems. |
| **Role Physical** | Limitations in usual role activities due to physical health problems. |
| **Bodily Pain** | Perception of pain in daily living |
| **General Mental Health** | Psychological distress and well-being |
| **Role Emotional** | Limitations in usual role activities due to emotional problems |
| **Vitality** | Energy and fatigue in daily living |
| **General Health** | Overall perception of physical health |

**Supplementary Table 2.**
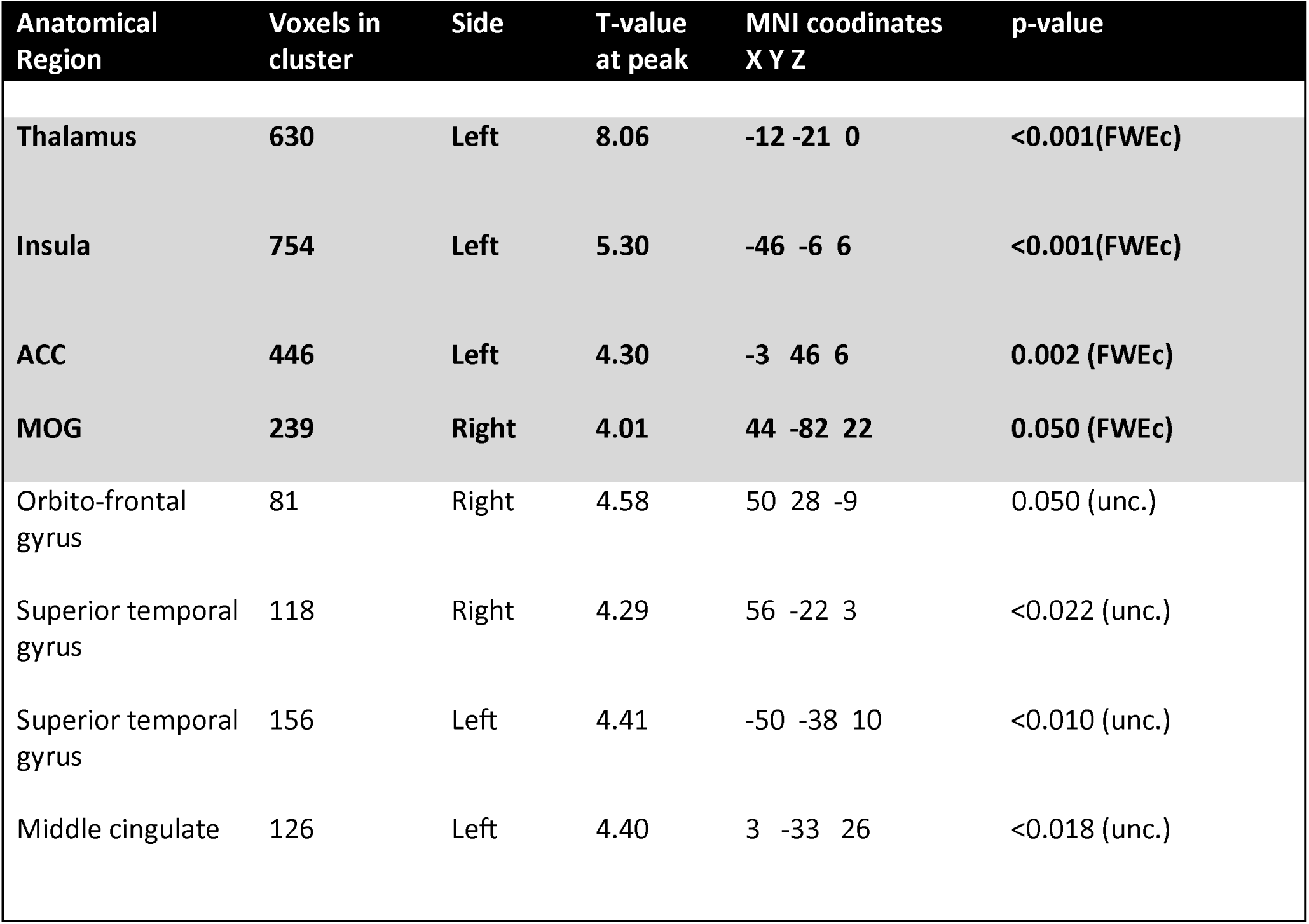
GMV Decreases per cluster. Gray-marked areas indicate significant FWE-corrected clusters, while unmarked areas represent uncorrected significant clusters. FWE cluster-level correction was applied with a threshold of k = 239.

GMV Decreases per cluster
| Anatomical Region | Voxels in cluster | Side | T-value at peak | MNI coordinates X Y Z | p-value |
| --- | --- | --- | --- | --- | --- |
| Thalamus | 630 | Left | 8.06 | -12 -21 0 | <0.001(FWEc) |
| Insula | 754 | Left | 5.30 | -46 -6 6 | <0.001(FWEc) |
| ACC | 446 | Left | 4.30 | -3 46 6 | 0.002 (FWEc) |
| MOG | 239 | Right | 4.01 | 44 -82 22 | 0.050 (FWEc) |
| Orbito-frontal gyrus | 81 | Right | 4.58 | 50 28 -9 | 0.050 (unc.) |
| Superior temporal gyrus | 118 | Right | 4.29 | 56 -22 3 | <0.022 (unc.) |
| Superior temporal gyrus | 156 | Left | 4.41 | -50 -38 10 | <0.010 (unc.) |
| Middle cingulate | 126 | Left | 4.40 | 3 -33 26 | <0.018 (unc.) |

**Supplementary Table 2.**
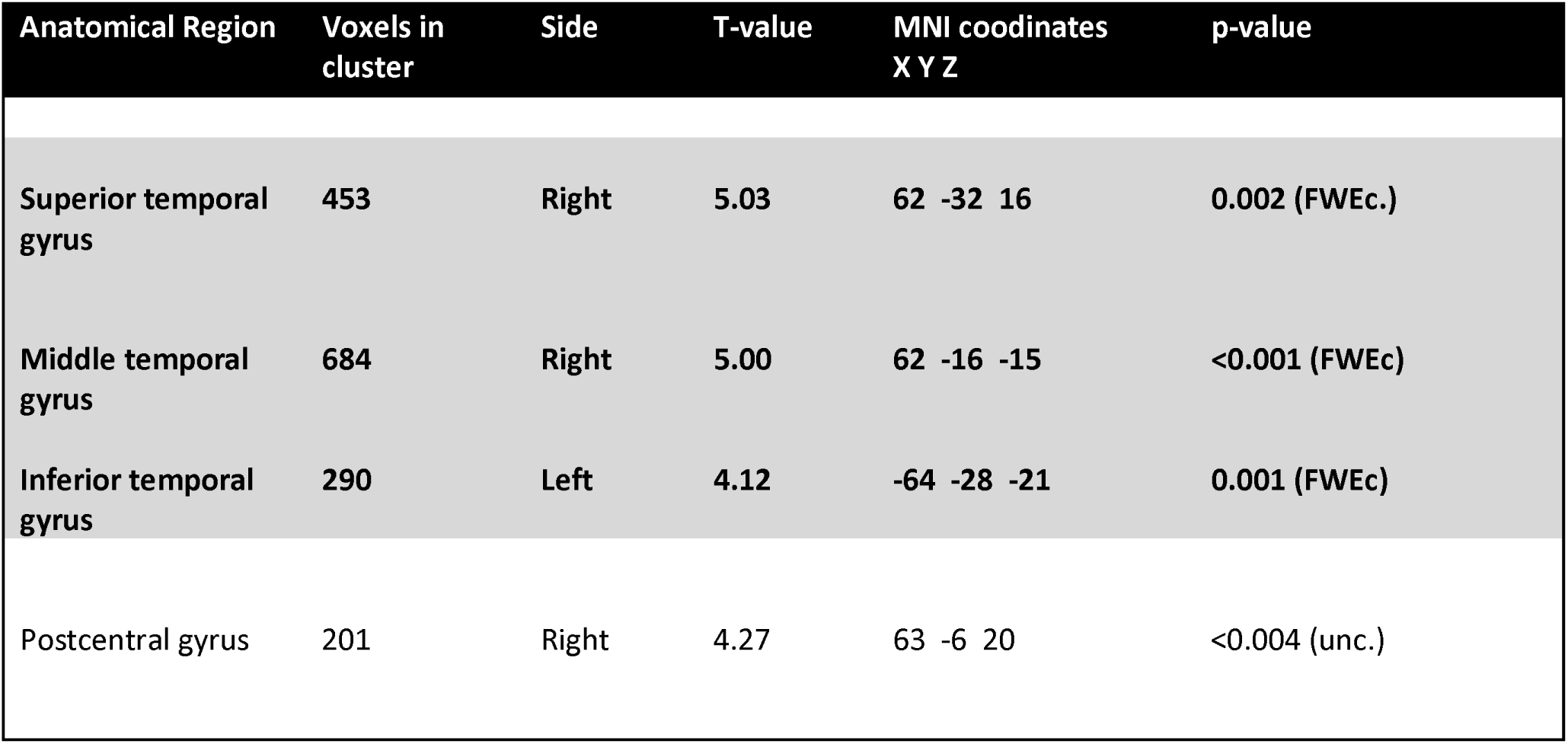
GMV Increases per cluster. Gray-marked areas indicate significant FWE-corrected clusters, while unmarked areas represent uncorrected significant clusters. FWE cluster-level correction was applied with a threshold of k = 290.

## Financial Disclosures of all authors (for the preceding 12 months)

Full Financial Disclosures of all Authors for the Past Year: Information concerning all sources of financial support and funding for the preceding twelve months, regardless of relationship to current manuscript, must be submitted with the following categories suggested.

List sources or “none”.

Stock Ownership in medically-related fields

Intellectual Property Rights

Consultancies

Expert Testimony

Advisory Boards

Employment

Partnerships

Inventions

Contracts

Honoraria

Royalties

Patents

Grants

Other

## Notes

### Competing Interest Statement

The authors have declared no competing interest.

