## Supplementary data for "Beyond the CTCT: Remote Structural Changes After VIM-MRgFUS in Essential Tremor"

**Supplementary material**

**Supplementary Table 1**

**Used clinical scales and descriptions**

| FTM  Domain | Description |
| --- | --- |
| Part A  Part B  Part C | A quantitative visual rating of tremor severity in different body parts (e.g., face, upper extremities, lower extremities) with respect of tremor type (e.g. postural or intention tremor)  An evaluation of the patient's ability to perform tasks requiring steady line drawing and writing.  A subjective assessment of limitations in activities of daily living (ADL; e.g., eating, drinking, hygiene). |
| **SF-36**  **Domain** | **Description** |
| **Physical Functioning**  **Social Functioning** | Limitations in physical activities due to health problems  Limitations in social activities due to physical or emotional problems. |
| **Role Physical**  **Bodily Pain**  **General Mental Health**  **Role Emotional**  **Vitality**  **General Health** | Limitations in usual role activities due to physical health problems.  Perception of pain in daily living  Psychological distress and well-being  Limitations in usual role activities due to emotional problems  Energy and fatigue in daily living  Overall perception of physical health |

**Supplementary Table 2**

**GMV Decreases per Cluster**

| Anatomical Region | Voxels in cluster | Side | T-value at peak | MNI coodinates  X Y Z | p-value |
| --- | --- | --- | --- | --- | --- |
| **Thalamus** | **630** | **Left** | **8.06** | **-12 -21 0** | **<0.001(FWEc)** |
| **Insula** | **754** | **Left** | **5.30** | **-46 -6 6** | **<0.001(FWEc)** |
| **ACC** | **446** | **Left** | **4.30** | **-3 46 6** | **0.002 (FWEc)** |
| **MOG** | **239** | **Right** | **4**.**01** | **44 -82 22** | **0.050 (FWEc)** |
| Orbito-frontal gyrus | 81 | Right | 4.58 | 50 28 -9 | 0.050 (unc.) |
| Superior temporal gyrus  Superior temporal gyrus  Middle cingulate | 118  156  126 | Right  Left  Left | 4.29  4.41  4.40 | 56 -22 3  -50 -38 10  3 -33 26 | <0.022 (unc.)  <0.010 (unc.)  <0.018 (unc.) |

*Table legend.* Gray-marked areas indicate significant FWE-corrected clusters, while unmarked areas represent uncorrected significant clusters. FWE cluster-level correction was applied with a threshold of k = 239.

**Supplementary Table 3**

**GMV Increases per Cluster**

| Anatomical Region | Voxels in cluster | Side | T-value | MNI coodinates  X Y Z | p-value |
| --- | --- | --- | --- | --- | --- |
| **Superior temporal gyrus** | **453** | **Right** | **5.03** | **62 -32 16** | **0.002 (FWEc.)** |
| **Middle temporal gyrus** | **684** | **Right** | **5.00** | **62 -16 -15** | **<0.001 (FWEc)** |
| **Inferior temporal gyrus** | **290** | **Left** | **4.12** | **-64 -28 -21** | **0.001 (FWEc)** |
| Postcentral gyrus | 201 | Right | 4.27 | 63 -6 20 | <0.004 (unc.) |

*Table legend.* Gray-marked areas indicate significant FWE-corrected clusters, while unmarked areas represent uncorrected significant clusters. FWE cluster-level correction was applied with a threshold of k = 290.
